# CD300e modulates metabolic programs in adipose tissue macrophages during obesity

**DOI:** 10.1101/2025.07.02.662518

**Authors:** Sara Coletta, Simone Pizzini, Stefania Vassallo, Elisabetta Trevellin, Annica Barizza, Sofia Giacometti, Gabriele Sales, Mattia Laffranchi, Silvano Sozzani, Marta Murgia, Agnese De Mario, Cristina Mammucari, Fabrizia Carli, Amalia Gastaldelli, Roberto Vettor, Marina de Bernard

**Affiliations:** Department of Biology, University of Padova, Padova, Italy; Department of Medicine (DIMED), University of Padova, Padova, Italy; Department of Molecular Medicine, Sapienza University of Rome, Laboratory Affiliated to Istituto Pasteur-Fondazione Cenci Bolognetti, Rome, Italy; Department of Biomedical Sciences, University of Padova, Padova, Italy; Department of Proteomics and Signal Transduction, Max-Planck Institute of Biochemistry, Martinsried, Germany; Cardiometabolic Risk Unit, Institute of Clinical Physiology, National Research Council-CNR, Pisa, Italy; Center for Metabolic and Nutrition Related Diseases - Humanitas Research Hospital Rozzano, Milano, Italy

**Keywords:** CD300e, obesity, macrophages, mitochondria, metabolism

## Abstract

Previous studies in monozygotic twins discordant for body mass index (BMI) revealed that obese individuals exhibit elevated expression of the immune receptor CD300e in white adipose tissue (WAT). Notably, CD300e levels decreased following weight loss, implicating its involvement in adipose tissue remodeling and metabolic regulation.

To elucidate the functional role of CD300e, we employed a Cd300e knockout (*Cd300e⁻/⁻*) mouse model subjected to a high-fat diet (HFD). Our findings demonstrate that CD300e deficiency exacerbates obesity-associated metabolic dysfunction, including increased weight gain, adipocyte hypertrophy, hepatic steatosis, and impaired glucose and insulin sensitivity. Adipose tissue macrophages (ATMs) lacking CD300e displayed reduced lipid and glucose uptake, alongside diminished mitochondrial respiration—a phenotype consistent with a broader metabolic impairment, as evidenced by proteomic profiling. These metabolic deficits were genotype-dependent and persisted after 16 weeks of HFD. Concurrently, adipocytes from *Cd300e⁻/⁻* mice exhibited enhanced lipogenesis and attenuated lipolysis. Remarkably, the impaired metabolic fitness exhibited by *Cd300e⁻/⁻* mouse macrophages was recapitulated in human cells upon gene silencing.

Collectively, these results establish CD300e as essential for ATM metabolic activation, positioning it as a key regulator of adipose tissue homeostasis and a critical mediator of obesity-induced metabolic dysfunction. Given its pivotal role, CD300e emerges as a promising therapeutic target for modulating adipose tissue function and improving metabolic health in obesity.

## Introduction

Obesity is recognized as a multifactorial chronic disease in which structural and functional modifications of adipose tissue (AT) have been recognized as the major trigger for the development of its complications^1–4^.

When the metabolic microenvironment changes due to obesity or excess lipids, quantitative and functional changes of innate immune cells occur along with the secretion of adipokines and pro-inflammatory cytokines that modulate the local microenvironment, promote insulin resistance, induce hyperglycaemia, and activate inflammatory signalling pathways. Many immune cells in AT have been described as participating in obesity-associated inflammation, among them macrophages constitute a large proportion.

The advances in single-cell RNA sequencing technologies have unveiled the remarkable heterogeneity of adipose tissue macrophages (ATMs), the most abundant immune cells in expanded white adipose tissue (WAT). Rather than conforming to the traditional M1/M2 paradigm, ATMs exhibit a diverse range of phenotypes and functions that are significantly altered in obesity^5,6^. This phenotypic diversity has been observed in both humans and mouse models, although the relative abundance of ATM subtypes may vary across species^7,8^. Mounting evidence highlights the critical crosstalk between adipocytes and ATMs as a key regulator of adipose tissue homeostasis, inflammation, and obesity-induced metabolic dysfunction^9^.

In obese conditions, ATMs accumulate in WAT and show transcriptional signatures enriched in pathways associated with inflammation, lipid uptake, lysosomal biogenesis and function, and lipid metabolism, including mitochondrial β-oxidation and respiration. They are commonly referred to as lipid-associated macrophages (LAM)^10^. Consistent with this molecular profile, LAMs exhibit a metabolically specialized yet immunologically activated phenotype, integrating lipid handling with inflammatory signaling as part of an adaptive response to metabolic stress¹⁰.

Under physiological conditions, adipocytes release fatty acids (FFA) and glycerol to support systemic energy demands, a process dependent on neutral lipases. Among these, adipose triglyceride lipase (ATGL) catalyzes the initial and rate-limiting step of lipolysis^11^. Once released, FFA can enter the bloodstream and taken up by peripheral tissues like liver, heart, and skeletal muscle. In addition, FFA can be locally engulfed by macrophages and processed through distinct metabolic pathways: β-oxidation for energy generation, esterification into triacylglycerols (TAG) for lipid storage, or cellular export^12^.

During the progression of obesity, the number of LAMs in adipose tissue increases due to both local proliferation and the infiltration of monocytes from the circulation^13^. Through their activity, these cells help buffer excess lipid–induced lipotoxicity and contribute to maintaining systemic homeostasis^10^. However, when lipid burden becomes excessive or chronic, the lipid-buffering capacity of LAMs may be overwhelmed, leading to a relative predominance of their inflammatory programs and contributing to adipose tissue inflammation and metabolic dysfunction^9,10^. Sustained inflammatory signaling within adipose tissue is a recognized driver of insulin resistance and systemic metabolic disease^14,15^.

CD300 family members comprise a group of paired activating and inhibitory immunoglobulin-like receptors (CD300a-h) that function as a lipid-sensing network^16^. They have been extensively studied for their role in the immune system and most of the family members are highly expressed within immune cells with little to no expression in other cell types^16–18^.

The immune receptor CD300e, expressed almost exclusively on myeloid cells^19,20^, has been detected in macrophages infiltrating the adipose tissue of patients with obesity^21^, and has recently garnered attention for its potential involvement in adipose tissue homeostasis. Notably, CD300e expression is upregulated in WAT from obese individuals and downregulated following weight loss^22–24^, indicating a tight association with metabolic status.

Given that adipose tissue expansion during obesity is accompanied by increased infiltration and phenotypic remodeling of macrophages, these observations raised the question of whether CD300e expression reflects a secondary consequence of macrophage accumulation in obese adipose tissue or, alternatively, whether CD300e expression by ATMs actively contributes to the regulation of adipose tissue homeostasis. Specifically, it remained unclear whether CD300e upregulation represents a maladaptive feature of obesity-associated inflammation or instead plays a protective role by modulating macrophage function and mitigating obesity-induced metabolic dysfunction.

To investigate this question, we used a Cd300e knockout mouse model to define the functional role of this immune receptor in the crosstalk between adipocytes and ATMs and its impact on obesity-associated adipose tissue remodelling and systemic glucose homeostasis.

## Results

### CD300e expressed by ATMs prevents the adipocyte hypertrophy induced by excessive energy intake

To investigate the functional role of CD300e in adipocyte-ATMs crosstalk, we employed a Cd300e knockout (KO) mouse model generated by the deletion of exon 2 of the *Cd300e* gene^25^ (Fig. 1A) and explored the impact of the gene deletion in adipose tissue remodelling and glucose homeostasis in diet-induced obesity.

**Figure 1.**
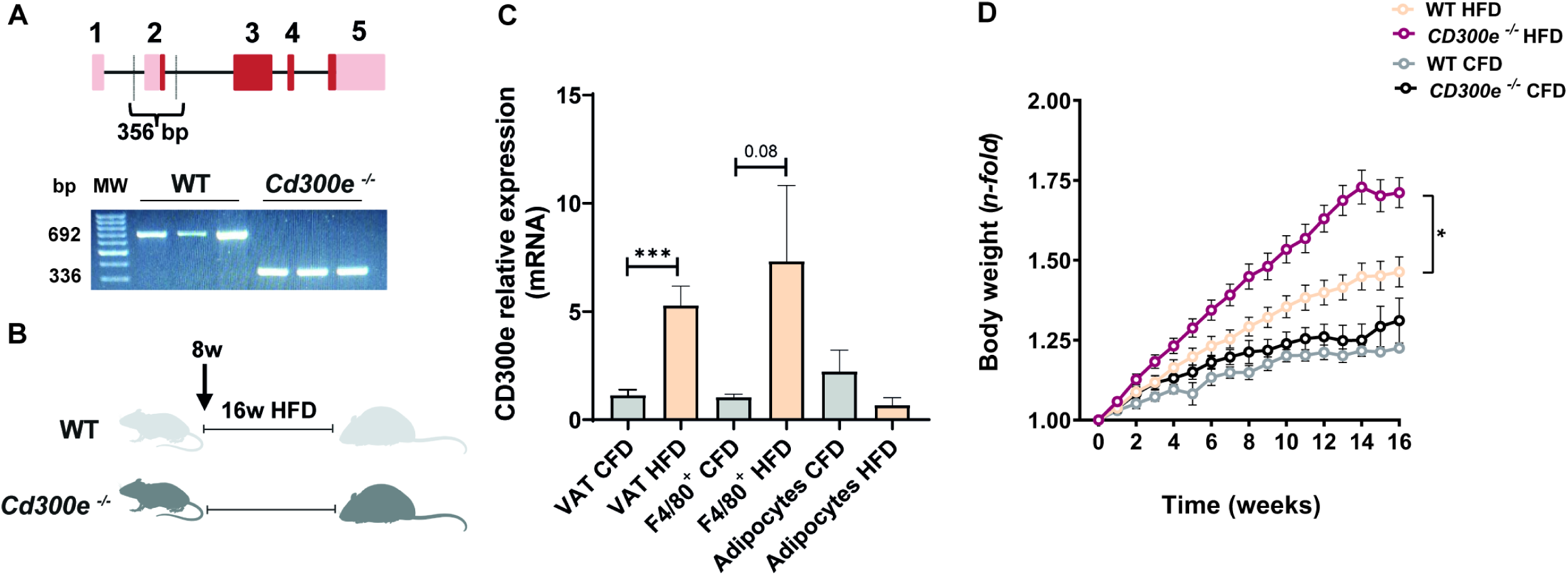
CD300e expression increases in ATMs during obesity and limits weight gain. (A) Upper panel: Schematic representation of the mouse *Cd300e* gene showing introns (black line) and exons (red boxes); the latter include untranslated regions (UTRs; light red) and protein-coding regions (dark red). *Cd300e^−/−^*mice carry a deletion of 356 bp encompassing exon 2, together with flanking intronic regions. Lower panel: Representative Cd300e genotyping results of F4/80^+^ cells isolated from VAT of WT and *Cd300e^−/−^*mice. PCR products were resolved by agarose gel electrophoresis. (B) Schematic representation of the diet-induced obesity model. (C) *Cd300e* mRNA expression in total VAT, F4/80^+^ cells and adipocytes, isolated from VAT of WT mice after 16 weeks of control chow diet (CFD) or high fat diet (HFD). mRNA levels were quantified by qRT-PCR and data were normalized to the endogenous reference gene 18S. Expression levels in HFD-fed mice are shown relative to CFD-fed mice. Data are presented as mean ± SEM (CFD, n=7; HFD, n=9). Significance was determined by Student’s t-test. ***, p< 0.001. (D) Body weight of WT and *Cd300e^−/−^* mice under CFD or HFD feeding. Data are shown as mean ± SEM (CFD, n=10 per genotype; HFD, n=22 per genotype). Statistical significance was determined by one-way ANOVA test. *, p< 0.05.

Before conducting a detailed investigation, we first assessed whether the upregulation of CD300e observed in adipose tissue of obese humans was also recapitulated in mice fed a HFD. WT mice of the C57BL/6 strain at the age of 8 weeks were fed a HFD ad libitum for 16 weeks (Fig.1B), while control WT mice were fed a CFD.

Fig. 1C shows that, like observations in humans, obese mice exhibited increased expression of *Cd300e* mRNA in epididymal visceral adipose tissue (VAT). Furthermore, in accordance with the almost exclusive expression of the immune receptor by myeloid cells, the diet-induced up-regulation was evidenced in ATMs (F4/80 positive cells) but not in adipocytes. These findings support the use of mice as a relevant model to study the role of the immune receptor in the context of obesity.

To investigated whether CD300e up regulation was part of a positive feedback loop activated by body mass gain to mitigate the harmful effects of overnutrition we utilized *Cd300e*-null mice subjected to a HFD and compared their phenotype to that of WT controls (Fig. 1B). There was no notable difference in body weight between WT and *CD300e^−/−^* mice under standard conditions (Fig. 1D). However, following HFD feeding, *Cd300e^−/−^* mice exhibited significantly greater body weight than their WT counterpart, with differences emerging as early as 6 weeks into the HFD feeding (Fig.1D). VAT and subcutaneous inguinal adipose tissue (SAT) were collected from all animals upon completion of the individual diet regimens. Macroscopic comparison of the VAT specimens indicated that *Cd300e*-null mice fed a HFD had a bigger fat mass depot than their WT counterparts subjected to the same diet (Fig. 2A, right panel). In accordance, while VAT weight was comparable between WT and *Cd300e^−/−^*mice fed a CFD, it was significantly increased in HFD-fed null mice compared to WT counterparts (Fig. 2A, left panel). H&E staining revealed considerable enlargement of adipocyte cell sizes in *Cd300e^−/−^*mice under HFD feeding, with quantification of the cross-sectional areas confirming the adipose tissue growth by cell hypertrophy (Fig. 2B).

**Figure 2.**
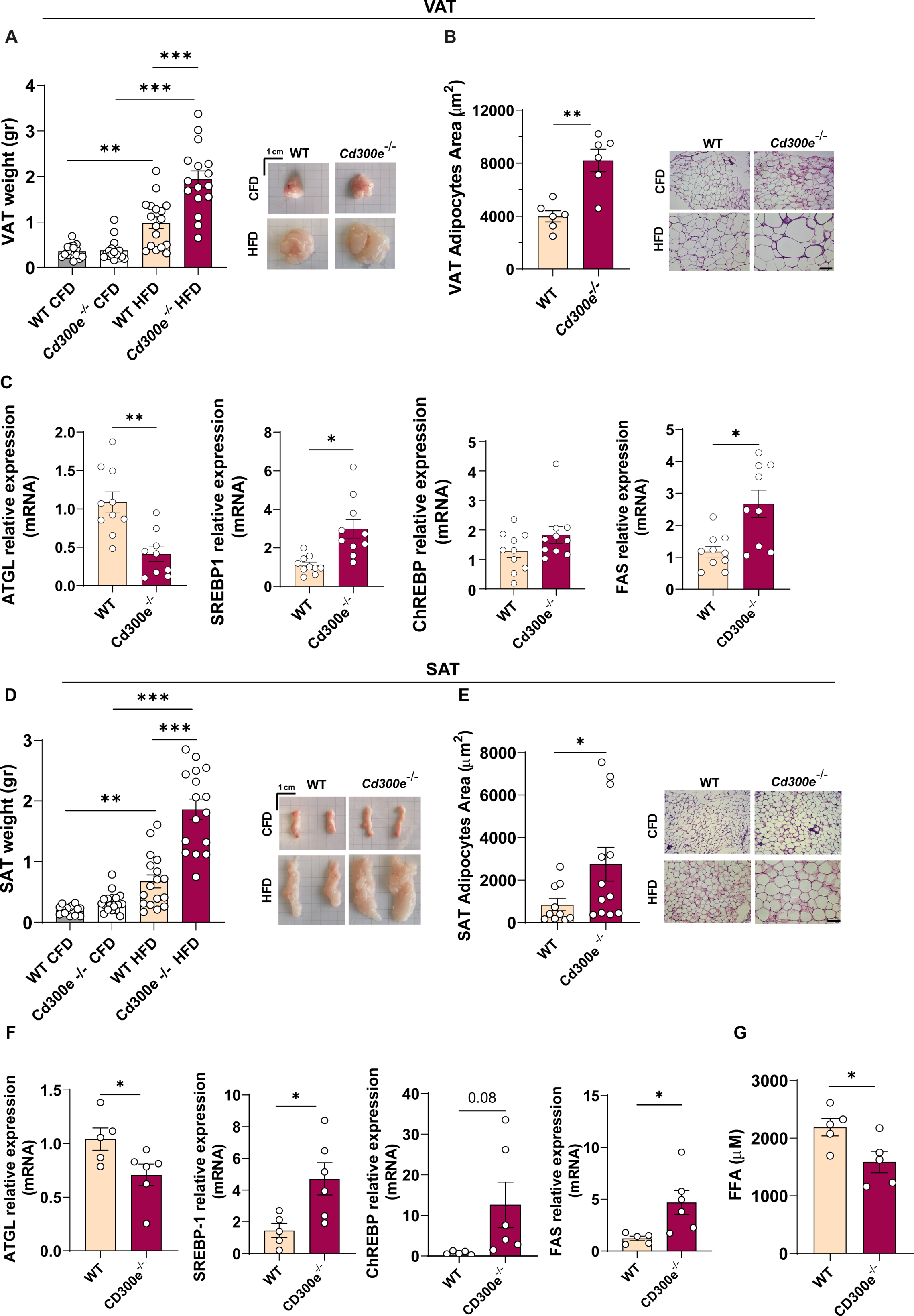
CD300e prevents the adipocyte hypertrophy induced by excessive energy intake. (A) and (D) VAT and SAT mass of WT and *Cd300e^−/−^* mice after 16 weeks of CFD or HFD feeding; data are shown as mean ± SEM (n=16/group). Significance was determined by one-way ANOVA test. **, p< 0.01; ***, p<0.001. Representative images of gross morphology of VAT and SAT from mice of the indicated genotype are reported. (B) and (E) Cross sectional area of VAT and SAT adipocytes of WT and *Cd300e^−/−^*mice after 16 weeks of HFD feeding (VAT: n=286 adipocytes from WT mice; n=280 adipocytes from *Cd300e^−/−^* mice; adipocytes derived from 6 mice per genotype; SAT: n=917 adipocytes from WT mice; n=750 adipocytes from *Cd300e^−/−^*mice; adipocytes derived from 10 WT mice and 12 *Cd300e^−/−^* mice). Significance was determined by Student’s t-test. *, p< 0.05; **, p< 0.01. Sections of VAT from WT and *Cd300e^−/−^* mice after CFD or HFD feeding were generated and stained with H&E. Scale bar = 100 μm. (C) and (F) mRNA expression of ATGL, SREBP1, ChREBP and FAS were evaluated by qRT-PCR on VAT and SAT of WT and *Cd300e^−/−^* mice after 16 weeks of HFD feeding. Data were normalized on 18S as endogenous reference genes and are expressed as 2^−ΔΔCt^ relative to WT mice. Data are expressed as mean ± SEM (VAT: WT, n=10; *Cd300e^−/−^*, n=9; SAT WT, n=5; *Cd300e^−/−^*, n=6). Statistical significance was determined by Student’s t test. *, p< 0.05; **, p< 0.01. (G) FFA were quantified on plasma of WT and *Cd300e^−/−^* mice after 16 weeks of HFD feeding. Data are expressed as mean of µM (n=5 per genotype). Statistical significance was determined by Student’s t test. *, p< 0.05.

Given the evidence that ATMs regulate lipid flux out of adipose tissue and that lipid-laden ATMs promote the production of yet unidentified antilipolytic factors^26^, we hypothesized that *Cd300e^−/−^*macrophages may impair adipocyte lipid catabolism, thereby contributing to adipocyte hypertrophy under nutrient-rich conditions.

To test this, we analyzed adipocytes from the VAT of obese animals and found that ATGL, the rate-limiting enzyme of triglyceride hydrolysis was significantly reduced in *Cd300e^−/−^* animals compared with WT controls (Fig. 2C). Consistent with this finding, adipocytes from KO mice exhibited a marked upregulation of the lipogenic program, with increased expression of SREBP-1, ChREBP, and FAS (Fig. 2C). These alterations were similarly observed in the SAT, Fig. 2D-F. Consistent with reduced adipocyte lipolysis, circulating levels of free fatty acids (FFA) were lower in KO mice compared to wild-type (WT) controls (Fig. 2G).

### CD300e prevents glucose dysmetabolism and liver steatosis during obesity

To determine the metabolic consequences of *Cd300e* ablation, we conducted GTT and ITT in both *Cd300e^−/−^* and WT mice on a CFD and HFD. In CFD mice, the deficiency of *Cd300e* did not alter glucose levels compared to WT, in both GTT and ITT (Fig. 3A and B). However, when HFD mice were challenged with an intraperitoneal glucose bolus, blood glucose levels were significantly higher in *Cd300e^−/−^* mice at 60, 90, and 120 minutes after the administration of the bolus compared to WT mice. Similarly, when HFD mice were injected with insulin, *Cd300e^−/−^*mice displayed higher blood glucose levels than WT mice at all time points, with significant differences at 15, 30, and 60 minutes after the hormone injection (Fig. 3A and, B). Furthermore, while the basal level of glucose, measured after a 5-hour fasting period, was comparable between WT and *Cd300e^−/−^* mice on CFD, *Cd300e^−/−^* mice exhibited significantly higher glucose concentrations compared to WT animals on HFD (Fig. 3C).

**Figure 3.**
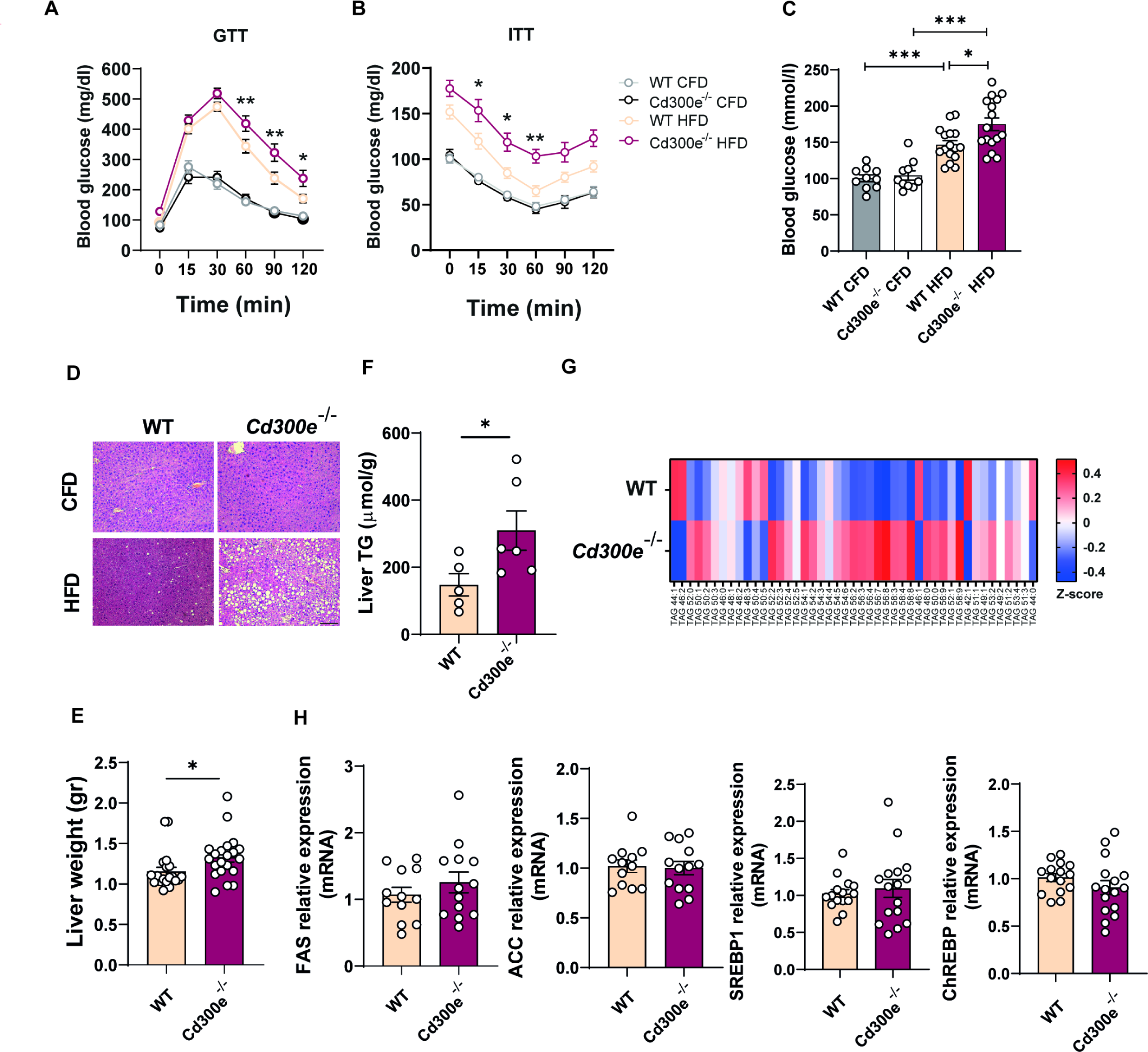
CD300e prevents glucose dysmetabolism and liver steatosis. (A and B) GTT and ITT of WT and *Cd300e^−/−^* mice after 16 weeks of CFD and HFD. Data are shown as mean ± SEM (CFD, n=12 per genotype; HFD, n=16 per genotype). Significance was determined by two-way ANOVA test. *, p< 0.05; **, p< 0.01. (C) Basal levels of glucose were measured in blood of 5-hours-fasted WT and *Cd300e^−/−^*mice at the end of the diet regimen. Data are shown as mean ± SEM (CFD, n=10 per genotype; HFD, n=16 per genotype). Significance was determined by one-way ANOVA test. *, p< 0.05; ***, p<0.001. (D) Sections of liver from WT and *Cd300e^−/−^* mice after 16 weeks of CFD or HFD were generated and stained with H&E. Scale bar = 100 μm. (E) Liver mass of WT and *Cd300e^−/−^* mice after 16 weeks of CFD or HFD feeding; data are shown as mean ± SEM (n=21 per group). Significance was determined by Student’s t test. *, p< 0.05. (F) Biochemical quantification of intrahepatic triglycerides (TAG) in liver of WT and *Cd300e^−/−^* mice after 16 weeks of HFD feeding. Data are shown as mean ± SEM (n=6 per genotype). Significance was determined by Student’s t-test. *, p< 0.05. (G) Heat map visualizing phenotypes of TAG lipidomes detected in liver of WT and *Cd300e^−/−^* mice after 16 weeks of HFD feeding. Data are expressed as mean of pmol/mg z-score (n=12 per genotype). (H) mRNA expression of FAS, ACC, SREBP1, and ChREBP were evaluated by qRT-PCR on Liver of WT and *Cd300e^−/−^* mice after 16 weeks of HFD feeding. Data were normalized on 18S as endogenous reference genes and are expressed as 2^−ΔΔCt^ relative to WT mice. Data are expressed as mean ± SEM (WT, n=12; *Cd300e^−/−^*, n=13).

Given the strong accumulation of lipids in the liver during obesity and the close association between adiposity and metabolic dysfunction-associated steatotic liver disease (MASLD) in humans^27^, we extended our analysis to this organ. The histological appearance of the liver was comparable between the two genotypes in CFD (Fig. 3D). However, *Cd300e^−/−^*mice fed a hypercaloric regimen exhibited more severe hepatic steatosis, which was accompanied by a significant increase in liver weight compared to WT mice (Fig. 3D and E). In accordance, total amount of TAG was higher in *Cd300e^−/−^*versus WT livers (Fig. 3F) and this was confirmed by lipidomic analysis that revealed that about 60% of major TAG species were increased in null mice (Fig. 3G). Notably, the expression of genes associated with *de novo* lipogenesis did not show any upregulation in *Cd300e^−/−^* mice after 16 weeks on HFD, compared to WT animals (Fig. 3H), in accordance with previous evidence that in advanced obesity-associated liver disease, hepatic steatosis can occur even in the absence of increased *de novo* lipogenesis, due to impaired VLDL assembly and secretion, which leads to intrahepatic lipid accumulation^28^.

Considering that CD300e is expressed almost exclusively by myeloid cells^29^, excluding neutrophils, we generated a conditional KO mouse lacking expression in the myeloid lineage to confirm that the observed effects were specifically due to the absence of CD300e in myeloid mononuclear cells. The results obtained using the conditional mutant closely mirrored those observed in the full KO mice in terms of weight gain, WAT remodelling, liver steatosis and impaired glucose metabolism (Fig. 4A-F).

**Figure 4.**
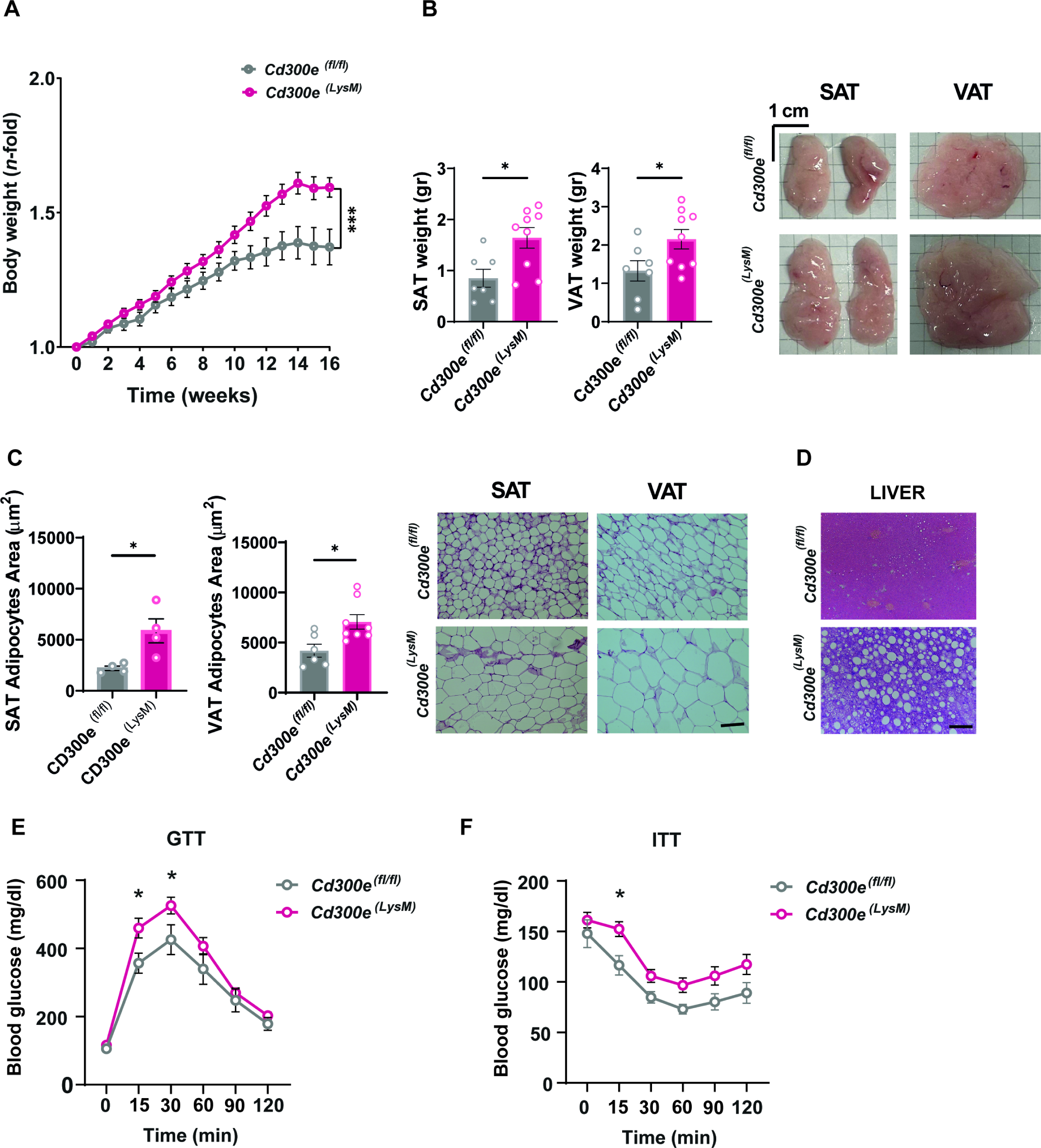
Characterization of myeloid-specific CD300e conditional knockout mice. (A) Bodyweight of *Cd300e^fl/fl^* and *Cd300e^LysM^*mice under HFD feeding. Data are shown as mean ± SEM (*Cd300e^fl/fl^*, n=7; *Cd300e^LysM^*, n=9). Significance was determined by Student’s t-test. ***, p< 0.001). (B) VAT and SAT mass of *Cd300e^fl/fl^* and *Cd300e^LysM^* mice after 16 weeks of HFD feeding; data are shown as mean ± SEM (*Cd300e^fl/fl^*, n=7; *Cd300e^LysM^,* n=9). Significance was determined by Student’s t-test. *, p< 0.05. Representative images of gross morphology of VAT and SAT from mice of the indicated genotype are reported. (C) Cross sectional area of VAT and SAT adipocytes of *Cd300e^fl/fl^* and *Cd300e^LysM^* mice after 16 weeks of HFD feeding (VAT: n=746 adipocytes from *Cd300e^fl/fl^*mice; n=922 adipocytes from *Cd300e^LysM^* mice; SAT: n=621 adipocytes from *Cd300e^fl/fl^* mice; n=433 adipocytes from *Cd300e^LysM^*mice. VAT adipocytes derived from 6 mice for *Cd300e^fl/fl^* and from 8 mice for *Cd300e^LysM^*, SAT adipocytes derived from 3 mice/genotype). Significance was determined by Student’s t-test. *, p< 0.05. Sections of VAT from *Cd300e^fl/fl^* and *Cd300e^LysM^* mice after HFD feeding were generated and stained with H&E. Scale bar = 100 μm. (D) Sections of liver from *Cd300e^fl/fl^*and *Cd300e^LysM^* mice after HFD feeding were generated and stained with H&E. Scale bar = 100 μm. (E-F) GTT and ITT of *Cd300e^fl/fl^* and *Cd300e^LysM^*mice after 16 weeks of HFD. Data are shown as mean ± SEM (*Cd300e^fl/fl^*, n=7; *Cd300e^LysM^*, n=8). Significance was determined by two-way ANOVA test. *, p< 0.05.

### *Cd300e^−/−^* mice lack significant adipose tissue inflammatory response

Adipose tissue inflammation contributes to obesity-related metabolic complications such as insulin resistance and type 2 diabetes^30^. While macrophage accumulation in WAT is a hallmark of this process^6,10,31^, T cells also undergo quantitative and phenotypic changes in obesity and contribute to metabolic disfunction and systemic inflammation^32^.

Therefore, we investigated whether the impaired glucose metabolism observed in *Cd300e^−/−^*mice might be accompanied by exacerbated adipose tissue inflammation compared to WT animals, as might be expected given the extensive tissue remodelling in the absence of the gene.

Consistent with published findings, we confirmed that diet-induced obesity in WT mice led to an increased proportion of macrophages (F4/80^+^ cells), as well as CD4⁺ and CD8⁺ T cells in VAT, compared to chow-fed controls (data not shown). In *Cd300e^−/−^* mice, the percentage of these cells raised further (Fig. 5A). However, the local pro-inflammatory cytokines IL-6, TNF-α, and IL-1β did not exhibit any relevant difference in terms of up-regulation between the two genotypes, with the only exception of TNF-α, whose levels showed a slight increase at both mRNA and protein levels, (Fig. 5B and C). Notably, no differences were observed in blood mononuclear cell composition between WT and *Cd300e^−/−^*mice, irrespective of the diet regimen (Supplementary Fig. 1). Collectively, these data argue against a substantial contribution of inflammation to the worsened dysmetabolism associated with the extensive adipose tissue remodelling observed in the absence of the immune receptor.

**Figure 5.**
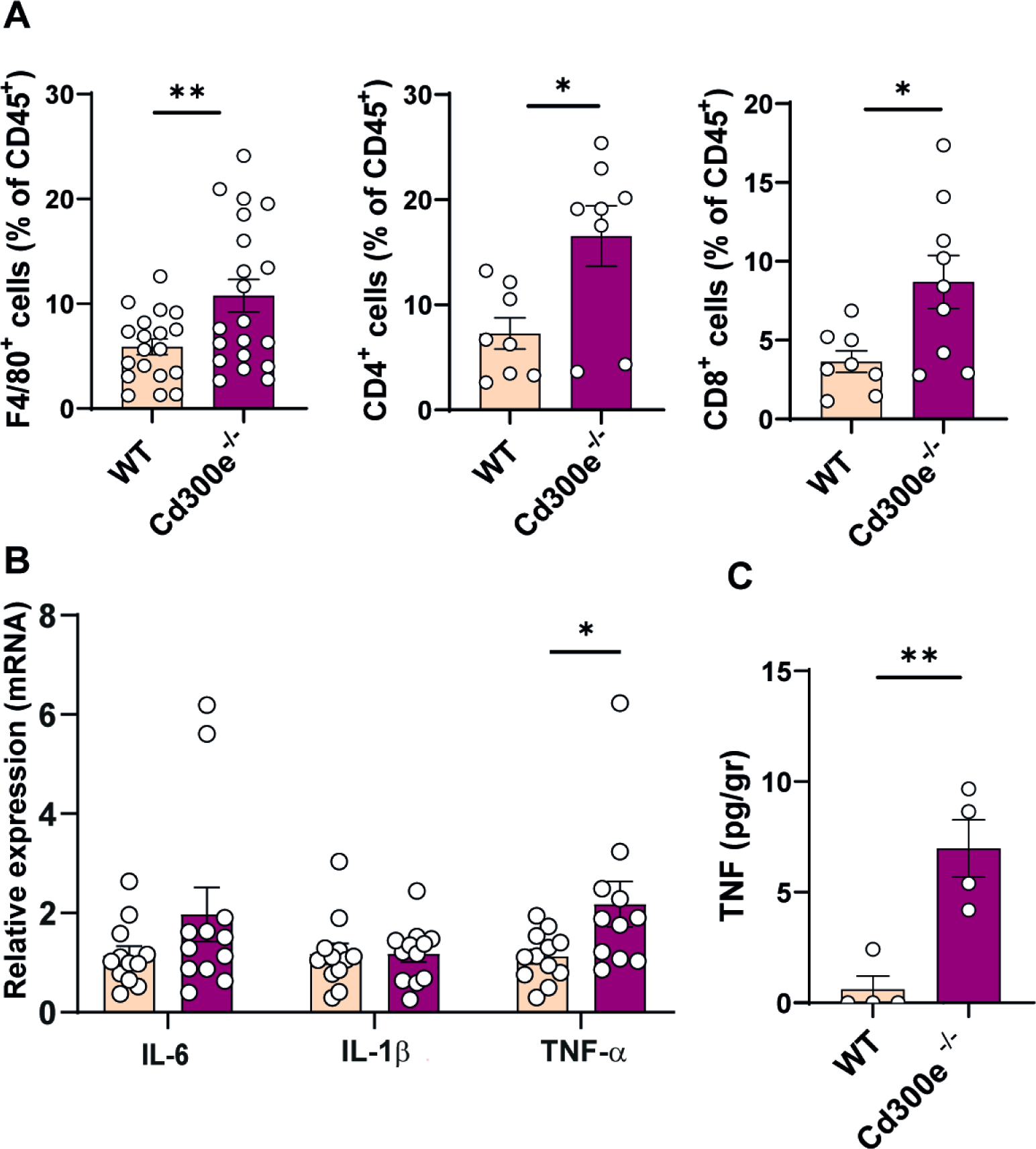
*Cd300e^−/−^* obese mice do not develop significant AT inflammation. (A) F4/80⁺ cells, CD4⁺ and CD8⁺ T lymphocytes (gated on CD45⁺ cells) were analyzed by flow cytometry in VAT infiltrates from WT and *Cd300e^−/−^*mice after 16 weeks of chow or HFD feeding. Data are shown as mean ± SEM (F4/80⁺ cells, n=15 per genotype; CD4⁺ and CD8⁺ T cells, n=8per genotype). Significance was determined by Student’s t test. *, p< 0.05. (B) mRNA expression of IL-6, IL-1β and TNF-α was evaluated by qRT-PCR on VAT of WT and *Cd300e^−/−^* mice after 16 weeks of HFD feeding. Data were normalized on 18S as endogenous reference genes and are expressed as 2^−ΔΔCt^ relative to WT mice. Data are expressed as mean ± SEM (VAT, n=12 per genotype). Statistical significance was determined by Student’s t test. P-values near the threshold for statistical significance are reported in the figure. (C) TNF-α was quantified on VAT digestion of WT and *Cd300e^−/−^* mice after 16 weeks of HFD feeding. Values are expressed as mean pg per gram of VAT tissue (n=4 per genotype). Statistical significance was determined by Student’s t test. *, p< 0.05.

### *Cd300e^−/−^* ATMs exhibit a profound metabolic impairment

To better characterize the phenotype of *Cd300e*^−/−^ ATMs, we isolated macrophages as F4/80^+^ cells from the VAT of 8-week-old WT or *Cd300e^−/−^* mice maintained on a control fat diet (T0) or subjected to HFD for further 16 weeks (HFD). These cells were subjected to mass spectrometry-based proteomics, focusing downstream analyses on the metabolic pathways where our experimental results pointed to specific alterations (Supplementary Fig. 2A-B, Supplementary Table S1).

Given the predominant role of ATMs metabolism in the AT homeostasis, we directed the protein abundance analysis on target proteins involved in pathways of lipid, glucose metabolism and energy production using the proDA tool^33^, testing genotype-dependent differences under both dietary conditions (Supplementary Fig. 2C and Supplementary Table S1).

Gene ontology analysis of enriched proteins according to ATM genotype at T0 or under HFD conditions revealed that proteins upregulated in *Cd300e* WT ATMs were predominantly associated with lipid metabolism, aerobic respiration and processes involved in the mitochondria-cell interactions (Fig. 6A). In contrast, *Cd300e^−/−^* ATMs were enriched for pathways related to inflammation and autophagy (Fig. 6A). As observed in the PCA (Supplementary Fig. 2B) diet *per se* emerged as a major driver of ATM proteome remodelling, irrespective of genotype. Therefore, to dissect how dietary conditions influence the differences induced by Cd300e ablation in ATMs, we performed a protein abundance analysis including diet as an interaction factor. Notably, even after accounting for dietary influence, ATMs from *Cd300e^−/−^* mice retained signatures of dysregulated lipid handling, an altered metabolic profile, and an overall higher inflammatory status (Fig. 6A).

**Figure 6.**
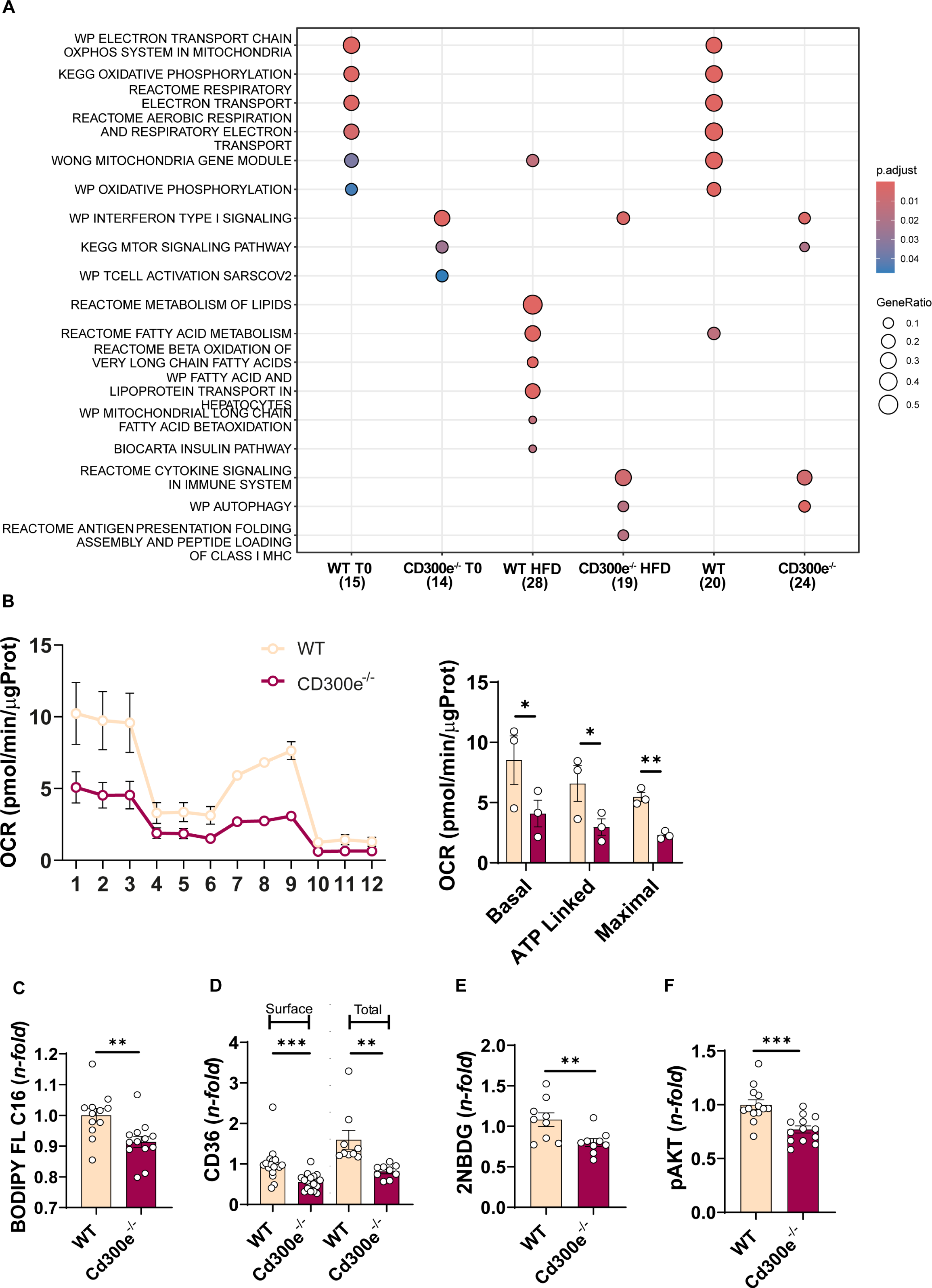
CD300e controls mitochondrial respiration and metabolic substrate uptake in macrophages. (A) Dot plot of gene ontology terms associated with proteins enriched by genotype highlighting the metabolic dysfunction of *C 300e*^−/−^ ATM. The total number of enriched proteins identified per condition is indicated. (B) OCR was measure in MME obtained from WT and *Cd300e^−/−^* mice. To calculate basal and maximal respiration, non-mitochondrial O2 consumption was subtracted from absolute values. ATP-linked respiration was calculated as the difference between basal and oligomycin insensitive O2 consumption. Data were normalized by protein content. Left panel: representative traces. Right panel: OCR quantification. Data are shown as mean ± SEM (n=3/ per genotype). Statistical significance was determined by Student’s t test. *, p< 0.05; **, p< 0.01. (C and E) The uptake of BODIPY FL C16 and 2-NBDG by MME from WT and *Cd300e*⁻/⁻ mice were quantified by flow cytometry. Data are expressed as mean ± SEM (Bodipy FL C16, n=13 per genotype; 2-NBDG, n=9 per genotype). Statistical significance was determined by Student’s t test. **, p< 0.01. (D) CD36 surface and total expression were analyzed by flow cytometry in MMe from WT and *Cd300e^−/−^* mice. Data are shown as mean ± SEM (surface, n=17 per genotype; total, n=9 per genotype). Statistical significance was determined by Student’s t test. **, p< 0.01; ***, p< 0.001. (F) Phosphorylation of AKT was analyzed by flow cytometry in MMe from WT and *Cd300e^−/−^* mice. Data are shown as mean ± SEM (n=13 per genotype). Significance was determined by Student’s t test. ***, p< 0.001.

Guided by proteomic evidence indicating impaired energy metabolism and compromised lipid handling in *Cd300e* null ATMs, we next sought functional validation of these alterations. The observation that ATMs isolated from the VAT of CFD-fed null mice already exhibited a metabolically hypoactive proteomic profile, even prior to HFD exposure, suggested that the metabolic defects were intrinsic to these cells rather than secondary to obesity, making them a suitable system to investigate the role of ATMs in adipose tissue remodelling while minimizing confounding effects associated with HFD-induced obesity.

To better characterize their bioenergetic and nutrient-handling properties in a controlled experimental setting, we employed bone marrow-derived monocytes isolated from null mice prior to initiation of the HFD regimen and differentiated them toward an MMe phenotype, which closely recapitulates the metabolically activated state of AT macrophages^34,35^. Using this *in vitro* model, we first assessed macrophage metabolism by performing extracellular flux analysis with the Seahorse platform. Compared with WT cells, *Cd300e^−/−^* MMe displayed significantly reduced basal mitochondrial respiration, ATP-linked respiration, and maximal respiratory capacity (Fig. 6B). These findings indicate that loss of the receptor markedly impairs mitochondrial respiration, underscoring its essential role in sustaining mitochondrial activity. Consistent with the impaired bioenergetic profile, receptor-deficient macrophages exhibited a reduced capacity to acquire metabolic substrates. Long-chain fatty acid uptake, assessed using BODIPY FL C16^36^, was significantly decreased in *Cd300e^−/−^*MMe (Fig. 6C). In parallel, flow cytometric analysis revealed reduced abundance of CD36, the fatty acid translocase that facilitates the uptake of long-chain fatty acids^37,38^, both at the cell surface and at the total cellular level following permeabilization (Fig. 6D). The concordant reduction of surface and total CD36 argues against a trafficking defect alone and instead points to diminished overall CD36 expression and/or stability. Finally, glucose uptake measured using the fluorescent analogue 2-NBDG was also markedly reduced in *Cd300e^−/−^* MMe (Fig. 6E). Together, these findings indicate a broad defect in nutrient acquisition, coherent with a metabolically hypoactive state.

Consistent with the impaired cellular energy phenotype, receptor-deficient macrophages exhibited reduced phosphorylation of AKT (Fig. 7F), a central node in nutrient-responsive signaling that promotes glucose uptake through translocation of glucose transporters to the cell surface^39^ and supports metabolic gene programs^40^.

**Figure 7.**
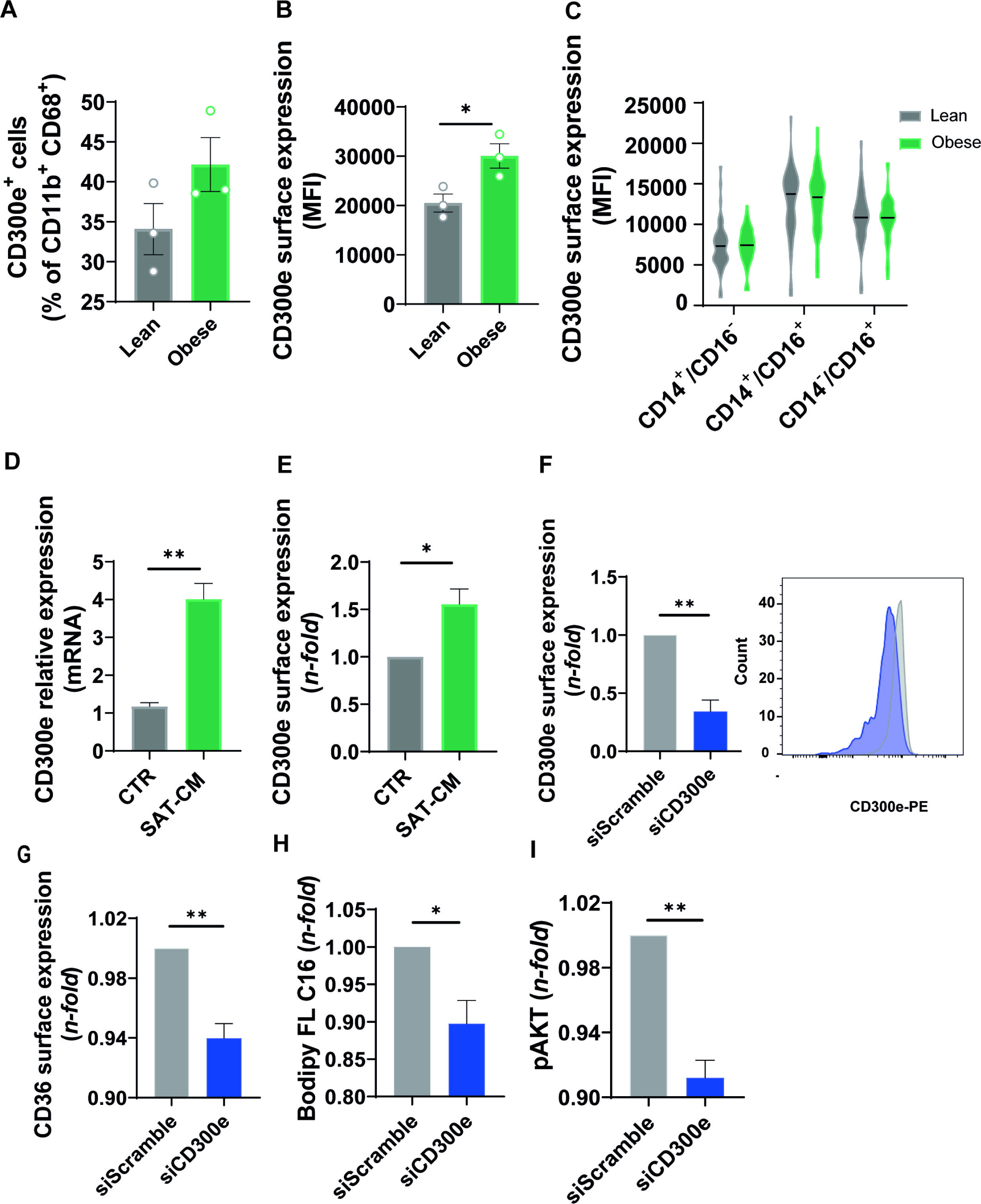
CD300e upregulation in ATMs regulates nutrient uptake and promotes AKT signaling in obesity. (A) Flow cytometry analysis of CD300e⁺ cells, gated on macrophages (CD11b⁺CD68⁺), in SVF from SAT of lean and obese subjects. Data are shown as mean ± SEM (n=3). (B) Surface expression of CD300e on macrophages (CD11b⁺CD68^+^ cells) infiltrating SVF of lean and obese SAT. Data are expressed as mean fluorescence intensity (MFI) ± SEM (n=3). (C) Violin plot of surface expression of CD300e on peripheral blood circulating monocytes of lean and obese patients. Circulating monocytes have been identified as classical (CD14^+^CD16^−^), intermediate (CD14^+^CD16^+^) and non-classical (CD14^−^CD16^+^). (lean, n=35; obese, n=43). (D) Expression of CD300e mRNA on freshly isolated monocytes cultured for 24 hours with conditioned medium derived from the SVF of obese SAT (SAT-CM). Data were normalized on β-actin as endogenous reference genes and are expressed as 2^−ΔΔCt^ relative to untreated monocytes (CTR). Data are expressed as mean ± SEM (SAT donors, n=4). (E) CD300e surface expression in monocytes stimulated as in (D). Expression levels of CD300e are represented as *n*-fold change relative to untreated monocytes set as 1 (n=4 independent experiments). Significance was determined by Student’s t-test. \**p*<0.05; **, p< 0.01. (F) left panel: surface expression of CD300e was monitored by flow cytometry in monocytes after 24 hours of silencing by siRNA. Expression levels of CD300e are expressed as *n*-fold change relative to untreated monocytes set as 1 (n=4 independently experiment). Significance was determined by Student’s t-test. **, p< 0.01. Right panel: representative histograms of CD300e surface expression in control and silenced monocytes. (G, H and I) CD36 surface expression, Bodipy FL C16 uptake and phosphorylation of AKT were analyzed by flow cytometry in monocytes after 24 hours of CD300e silencing. Data are shown as *n*-fold change relative to untreated monocytes set as 1 ± SEM (n=4). Significance was determined by Student’s t test. *, p< 0.05; **, p< 0.01.

### The adipose tissue microenvironment in human obesity induces CD300e expression during monocyte-to-macrophage differentiation, enabling regulation of nutrient uptake and AKT signaling

Based on prior evidence showing elevated CD300e mRNA levels in the adipose tissue of individuals with obesity, which decline after weight loss^23,24^, we sought to determine whether our murine findings had a human counterpart. First, we asked whether increased mRNA expression was associated with enhanced receptor surface expression on individual ATMs, rather than simply reflecting greater infiltration of these cells into the tissue. To this end, we isolated the stromal vascular fraction (SVF) from SAT samples collected from obese individuals, as well as from lean control subjects. ATMs were identified within the SVF as CD11b⁺CD68⁺ cells and subsequently analyzed for CD300e expression. As shown in Fig. 7A, a higher percentage of ATMs from formerly obese individuals expressed CD300e compared to those from lean subjects. Moreover, the mean fluorescence intensity (MFI) of CD300e was significantly greater in the obese group (Fig. 7B), suggesting enhanced receptor expression per cell.

Given that a large fraction of ATMs originate from infiltrating monocytes, and that monocyte recruitment increases with adipose tissue expansion^31,41,42^, we then assessed whether circulating monocytes from the two cohorts differed in CD300e expression. Consistent with previous findings, we confirmed that intermediate (CD14^+^CD16^+^) and non-classical monocytes (CD14^−^CD16^+^) exhibit slightly higher expression levels of CD300e compared to classical subset (CD14^+^CD16^−^) ^17^. However, no significant differences in CD300e surface expression were observed across any monocyte subtypes between subject with obesity and lean individuals (Fig. 7C). These findings suggest that CD300e upregulation occurs locally, likely induced by microenvironmental signals within the adipose tissue, during the differentiation of recruited monocytes into tissue-resident macrophages. In accordance, conditioned media collected from freshly isolated vascular fraction of obese SAT triggered the expression and the exposure of CD300e in monocytes (Fig. 7D and E).

Subsequently, we sought to determine whether reduced expression of CD300e in human myeloid cells would also result in impaired nutrient uptake and altered metabolic signaling, as observed in murine macrophages. To this end, we silenced CD300e in human monocytes, which not only represent the myeloid population with the highest receptor expression but are also more amenable to gene silencing than fully differentiated macrophages.

Using a cocktail of four siRNAs, we achieved a reduction in gene expression of more than 50% (Fig. 7F).

Remarkably, CD300e silencing resulted in reduced expression of CD36, consistent with a decrease in fatty acid uptake, and was accompanied by diminished AKT phosphorylation (Fig. 7G-I). Together, these findings indicate that reduced expression of CD300e in human myeloid cells is sufficient to impair metabolic fitness and nutrient sensing, thereby recapitulating key functional defects observed in *Cd300e^−/−^* murine macrophages.

## Discussion

In this study, we identify CD300e as a previously unrecognized regulator of adipose tissue macrophage (ATM) metabolic fitness and adipose tissue homeostasis in obesity. While CD300e expression is markedly increased in ATMs from obese mice and humans, our data demonstrate that this upregulation is not merely a phenotypic marker of adipose tissue inflammation, but reflects a functional requirement for maintaining macrophage metabolic competence in the obese adipose niche. Genetic ablation of Cd300e results in exacerbated diet-induced obesity, adipocyte hypertrophy, impaired systemic glucose tolerance and insulin sensitivity, and increased hepatic lipid accumulation. Importantly, these metabolic alterations are recapitulated in mice with myeloid-specific deletion of Cd300e, supporting a macrophage-intrinsic role for CD300e in controlling whole-body metabolic homeostasis. Notably, despite the worsening metabolic phenotype, loss of CD300e does not elicit a pronounced increase in classical inflammatory cytokines within adipose tissue, with the exception of a modest elevation in TNF-α. This dissociation between metabolic dysfunction and overt inflammatory burden suggests that additional mechanisms may be involved. Accordingly, we investigated whether CD300e supports macrophage-intrinsic metabolic programs. Consistent with this, proteomic profiling of Cd300e-deficient macrophages revealed broad alterations in pathways related to lipid handling and mitochondrial metabolism, indicating a central role for CD300e in sustaining ATM metabolic fitness. Functionally, CD300e loss was associated with reduced fatty acid and glucose uptake, diminished mitochondrial oxidative capacity, and impaired activation of AKT signaling, accompanied by decreased expression of the lipid transporter CD36. Together, these defects define a state of metabolic failure in CD300e-deficient macrophages, which likely compromises their ability to adapt to the lipid-rich obese adipose environment.

Emerging evidence suggests that metabolically competent ATMs play an active role in buffering excess lipids and coordinating adipose tissue remodeling during obesity. In this context, our findings support a model in which CD300e contributes to preserving ATM metabolic integrity, thereby indirectly sustaining adipocyte function. Consistent with this idea, adipose tissue from Cd300e-deficient mice displays reduced lipolytic capacity, increased expression of lipogenic markers, and lower circulating free fatty acid levels. While the precise macrophage-derived signals mediating this altered adipocyte program remain to be identified, our data indicate that impaired macrophage nutrient handling may disrupt macrophage–adipocyte communication, favoring maladaptive lipid storage and adipocyte hypertrophy.

Importantly, our mouse findings are mirrored in human obesity. CD300e expression is increased in ATMs isolated from adipose tissue of obese individuals and is induced in macrophages by adipose tissue–conditioned media, indicating that local adipose cues drive CD300e upregulation. Moreover, silencing CD300e in human macrophages recapitulates key features observed in mouse cells, including reduced CD36 expression, impaired AKT activation and diminished lipid uptake. These observations support the translational relevance of CD300e-dependent macrophage metabolic regulation in human adipose tissue.

Based on these results, we propose that CD300e functions as a component of a macrophage program that supports lipid uptake and metabolic adaptation in the obese adipose microenvironment. While members of the CD300 family have been implicated in lipid recognition and immune regulation, further studies will be required to define whether CD300e directly senses lipid ligands or instead modulates intracellular metabolic pathways downstream of adipose-derived signals. Nevertheless, our data identify CD300e as a critical node linking macrophage metabolic fitness to adipose tissue function.

In conclusion, this study reveals a non-canonical role for CD300e in maintaining ATM metabolic competence and preserving adipose tissue homeostasis during obesity. By supporting macrophage nutrient handling rather than suppressing overt inflammation, CD300e limits maladaptive adipose tissue remodeling and systemic metabolic dysfunction. Targeting macrophage metabolic fitness pathways such as CD300e may therefore represent a complementary therapeutic strategy to improve metabolic health in obesity-associated disease.

## Material and Methods

### Patients

Blood samples were collected from 43 obese patients (aged 18-60 years; with BMI > 40.0 kg/m^2^ or with BMI 35.0-39.9 kg/m^2^ with comorbidities) enrolled at the Center for the Study and the Integrated Treatment of Obesity of the University Hospital of Padova, and from 35 normal-weight subjects (aged 18-60 years; BMI <25 kg/m^2^) who served as healthy controls.

SAT was obtained from obese patients who underwent plastic surgery (abdominoplasty) after bariatric surgery-induced weight loss (n=3) and from lean individuals undergoing elective minor abdominal surgery (n=3). Exclusion criteria included: diagnosis of HIV and/or HCV; presence of any oncological disease at time of enrolment; chronic substance or alcohol abuse.

Anthropometric characteristics of patients are reported in Supplementary Table S2.

### Mice and diet

Experiments were conducted using WT, CD300e constitutive knockout (*Cd300e^−/−^*), floxed *Cd300e^(fl/fl)^,* and myeloid-specific knockout *Cd300e^(LysM)^* mice, all on a C57BL/6J background. Mice were housed in groups of 3 to 5 per cage at animal facility of the University of Padova under a 12 h light-dark cycle, with free access to water and to a standard chow diet (Mucedola S.r.l, Settimo Milanese, Italy). All experimental procedures were performed on male mice. Sample size was established according to previous studies ^43^. No criteria for animal exclusion were set.

Starting at age 8 weeks, mice randomly received chow diet as control fat diet (CFD) or 60% lard high-fat diet (HFD; Sniff Spezialditen GmbH, Soest, Germany) for 16 weeks. Body weight gain over the 16-week dietary intervention was the primary outcome measure and was recorded weekly. Mice were euthanized at 24 weeks of age by cervical dislocation.

### DNA extraction and genotyping

F4/80⁺ cells isolated from the VAT of WT or *Cd300e^−/−^* mice were homogenized in TRIzol reagent (Thermo Fisher Scientific, Waltham, MA, USA) by pipetting, and DNA was extracted according to the manufacturer’s protocol for genotyping. Following quantification using a NanoDrop Lite spectrophotometer (Thermo Fisher Scientific), 50 ng of DNA was used as template for PCR amplification, performed with DreamTaq DNA Polymerase (Thermo Fisher Scientific) according to the manufacturer’s instructions. A touchdown PCR protocol was employed to increase reaction specificity.

### Stromal vascular fraction (SVF) isolation and ATM purification

For human SVF isolation, SAT from 3 obese patients and 3 lean individuals was processed as previously described^21^.

For mouse SVF preparation, VAT and SAT fat pads were excised, finely minced, and incubated with 1 mg/ml collagenase type 2 (Gibco, Thermo Fisher Scientific) in DMEM/F-12 (Aurogene S.r.l., Rome, Italy) containing 10% fetal bovine serum (FBS, MicroGEM Ltd, Auckland, New Zealand), for 45 minutes at 37 °C in constant agitation. Cells were filtered through a 100 µm cell strainer and SVF was separated from mature adipocytes by centrifugation at 400*g* for 7 min. Supernatant and adipocyte fraction were removed and SVF was resuspended in DMEM/F-12 2% FBS and filtered through a 70 µm cell strainer. Erythrocytes were lysed in lysis buffer (BD Pharm Lyse; BD Biosciences, San Jose, CA, USA), washed in DMEM/F-12 2% FBS, and pellet was resuspended in DMEM/F-12 10% FBS for subsequent analysis.

For ATM isolation from SVF of VAT, after filtration through a 70 µm cell strainer, ATMs were isolated with anti-F4/80 Microbeads Ultrapure (Miltenyi Biotec, Bergisch Gladbach, Germany), following the manufacturer’s instructions.

### Bone marrow cells isolation and MME polarization

Murine Bone Marrow Derived Macrophages (BMDMs) were differentiated and polarized from bone marrow stem cells as previously described^34^. Briefly, bone marrow cells were obtained by flushing femurs and tibias with PBS 2% FBS. The cells were filtered through a 70 μm filter, and red blood cells were lysed via BD PharmLyse Lysing Buffer (BD Biosciences). To differentiate macrophages, bone marrow cells were seeded in Petri dishes in RPMI medium supplemented with 10% FBS, 100 U/ml penicillin, 100 µg/ml streptomycin, and 10% conditioned medium obtained from the L929 cell line. Macrophage differentiation was achieved 7 days after seeding. For MMe polarization, macrophages were treated with a combination of glucose (30 mM), insulin (10 nM), and palmitate (0.4 mM) for additional 24 hours.

### Preparation of human SAT stromal vascular fraction conditioned media

Stromal vascular fraction cells were isolated from subcutaneous adipose tissue of 4 patients undergoing plastic surgery and conditioned media were prepared as reported previously^44^. Briefly, cell suspension was seeded in DMEM/F12 supplemented with 10% FBS (5 × 10^5^ cells/well in 24-well plates). After 24 hours, cells were washed twice with sterile PBS (MicroGEM Ltd) and the medium was replaced with low-glucose DMEM (Gibco, Thermo Fisher Scientific) phenol red and FBS free. After 24 hours, conditioned medium was collected (SAT-CM) and instantly frozen in liquid nitrogen.

### Human peripheral blood monocyte isolation and treatment

Monocytes were purified from buffy coats of healthy donors by density gradient protocol, as described previously^45^.

To evaluate the effect of conditioned media obtained from stromal vascular fraction of SAT on CD300e expression, monocytes were plated at a density of 1.2 × 10^6^ per well in 24-well in 500 µl of RPMI (Gibco, Thermo Fisher Scientific) 10% FBS, 4 mM Hepes (Corning, NY, USA), 50 µg/ml of gentamycin (Gibco, Thermo Fisher Scientific), 50% SAT-CM. After 24 hours, cells were processed for subsequent analyses.

When monocytes were used for gene silencing experiments, cells were isolated using the Pan Monocyte Isolation Kit (Miltenyi Biotec) according to the manufacturer’s instructions.

For silencing experiments, a pool of four siRNAs targeting CD300e (QIAGEN, Hilden, Germany) was used in combination with the Cell Line Nucleofector™ Kit V (Lonza, Basel, Switzerland). Briefly, 7 × 10⁶ monocytes were resuspended in 100 μl of nucleofection buffer, and siRNAs were added at a final concentration of 150 nM each. AllStars Hs Negative Control siRNAs were used as a negative control.

The cell suspension was transferred to a nucleofection cuvette, and electroporation was performed using a Nucleofector IIb device (Lonza) with a program optimized for human monocytes.

Following nucleofection, cells were immediately recovered in RPMI medium without FBS, supplemented with 4 mM HEPES and 50 μg/ml gentamicin, and seeded in 24-well plates at a density of 1.1 × 10⁶ cells per well in 250 μl. Cells were incubated at 37°C in a humidified atmosphere containing 5% CO₂ for 1 hour, after which 250 μl of complete medium was added to achieve final concentrations of 20% FBS, 4 mM HEPES, 50 μg/ml gentamicin, and 100 ng/ml M-CSF (Miltenyi Biotec). After 24 hours, cells were harvested for subsequent analyses. SiRNA sequences are reported in Supplementary Table S3.

### Metabolic tolerance tests

One week before the conclusion of HFD protocol, glucose tolerance test (GTT) and insulin tolerance test (ITT) were conducted to determine HFD-induced alterations in glucose homeostasis and insulin sensitivity. For GTT, following an overnight fasting, mice underwent intra-peritoneal (i.p.) injection of glucose (2 g/kg, PanReac Applichem, Darmstadt, Germany). For ITT, following a 5-hours fasting, insulin 0.75 U/kg Humulin^®^ (Eli Lilly and Company, Indianapolis, IN, USA) was injected i.p. Glycemia was measured immediately before and at 15, 30, 60, 90, and 120 minutes after glucose or insulin administration. Glucose level was determined from tail blood using Accu-Chek Guide glucometer (Roche Diabetes Care, Mannheim, Germany).

### Tissue and blood collection from mice

Blood was collected by mandibular vein puncture immediately before sacrifice in EDTA-treated tubes. For plasma preparation, blood was centrifuged at 2,000g for 15 minutes at RT and the plasma transferred in a separate tube. After separation, plasma was stored at -80°C for further analysis. SAT, VAT, and liver were harvested and weighted. After rinsing in PBS, tissues were divided into small pieces (0.5 cm^3^). Pieces dedicated to RNA extraction or to lipidomic analysis were kept at - 80°C, while pieces intended for histological analysis were fixed in a 4% paraformaldehyde solution (Bio-Optica Milano S.p.A. (Milan, Italy) in PBS for 24 hours and then embedded in paraffin (BioOptica); tissues that were meant to stromal vascular fraction isolation and subsequent procedure were used immediately after the dissection.

### Tissue staining

Hematoxylin and eosin (H&E) staining was performed on 2-micron-thick formalin-fixed, paraffin-embedded VAT, SAT, and liver tissue sections. Bright field images were acquired with the DM6 microscope (Leica Microsystems, Wetzlar, Germany) using LasX software. VAT and SAT images were analyzed with Fiji (ImageJ) software to determine the area of adipocytes.

### Quantification of triglycerides, FFA and TNF-α

Hepatic triglycerides, plasma FFA and VAT TNF-α were measured using specific commercial kits, according to the manufacturers’ instructions. Hepatic triglyceride content was quantified using the Triglyceride Assay Kit (Abcam, Cambridge, UK). Plasma levels of FFA were determined with the FFA Assay kit (Abbexa Ltd, Cambridge, UK). TNF-α levels were measured in the supernatant of VAT digestion using a mouse TNF-α ELISA kit (Thermo Fisher Scientific).

### Lipid composition

Lipidomic analyses were performed in and liver tissues using liquid chromatography/quadrupole time-of-flight mass spectrometry (UHPLC -QTOF, 1290 Infinity-6545 Agilent Technology, Santa Clara, CA, USA) in positive ion ionization equipped with an Agilent ZORBAX Eclipse Plus C18 2.1 × 100 mm 1.8-Micron column at 50 °C. TAG were quantified using internal standard added to each sample, specifically TAG (15:0/15:0/15:0). Lipids were purchased from Avanti Polar Lipids (Alabaster, AL, USA) and Larodan (Solna, Sweden).

About 25 mg of liver tissues were homogenized using Precellys Evolution Homogenizer (Bertin Instruments, Frankfurt, Germany) (3 cycles of 30 seconds at 5800 rpm, with 20 seconds pause between each cycle, at 4 °C) and samples were centrifuged for 20 minutes at 12,900 rpm at 4 °C to remove proteins from samples. Lipid species were extracted with modified Folch method using 900 µl of chloroform:methanol (2:1) and 200 µl of water, centrifuged for 15 minutes at 12,900 rpm at 4 °C and down phases were dry under gentle nitrogen flux and resuspended in 200 µl of chloroform:methanol (1:1) solution.

### Flow Cytometry

#### Human subcutaneous SVF

SAT stromal vascular fraction was obtained as detailed above. Cells were resuspended in FACS buffer (PBS, 2% FBS) and incubated for 15 min at RT with Human TruStain FcX (Biolegend, San Diego, CA, USA) to block Fc receptors. 0.5 × 10^6^ cells were stained with the unconjugated monoclonal antibody anti-CD300e (clone UPH2, Thermo Fisher Scientific, MA5-44078) followed by a goat anti-mouse Alexa Fluor 647 antibody (Thermo Fisher Scientific, A21236), PE-Cy7-conjugated anti-CD45 (clone HI30, BD Biosciences, 557748) and BV785-conjugated anti-CD11b (clone M1/70, Biolegend, 101243). Zombie Aqua Fixable Viability Dye (Biolegend, 423101) was used to exclude dead cells. Cells were fixed with 3.6% formaldehyde and permeabilized with PermWash solution (1% FBS, 0.2% saponin in PBS). Cells were stained with PE-conjugated anti-CD68 monoclonal antibody (clone Y1/82A; eBioscience, Thermo Fisher Scientific, 12-0689-42), resuspended in FACS buffer, and acquired by LSR Fortessa X-20 Cell Analyzer (BD Biosciences). Data were analysed using FlowJo software, version 10.3 (Tree Star Inc., Ashland, OR, USA).

#### Human peripheral blood monocytes

150 ml of peripheral blood collected into EDTA-treated tubes were incubated for 5 minutes at RT with 3 ml of lysis buffer (BD Pharm Lyse, BD Biosciences) to lyse erythrocytes. Cells were washed with FACS buffer (PBS, 2% FBS) and incubated with Human TruStain FcX (Biolegend) for 15 minutes at RT to block Fc receptors. At the end of incubation cells were stained with the following monoclonal antibodies: PE-conjugated anti-CD300e (clone UP-H2, Biolegend, 339704), APC-conjugated anti-HLA-DR (clone L243, Biolegend, 307610), FITC-conjugated anti-CD3 (clone UCHT1, Biolegend, 300440), FITC-conjugated-CD19 (clone HIB19, eBioscience, 11-0199-42) and FITC-conjugated-CD56 (clone TULY56, eBioscience, 11-0566-42), PerCP/Cyanine5.5-conjugated anti-CD14 (clone 63D3, Biolegend, 367110) and BV785-conjugated anti-CD16 (clone 3G8, Biolegend, 302046). The BD Horizon™ fixable viability stain 780 (BD Biosciences) was used to exclude dead cells from the analysis. Cells were resuspended in FACS buffer and analysed by LSR Fortessa X-20 Cell Analyzer (BD Biosciences). The gating strategy adopted to analyse monocytes subsets was previously reported^46^ and illustrated in Supplementary Fig. 3. Data were analysed using FlowJo software, version 10.3 (Tree Star Inc., Ashland, OR, USA).

#### Human monocytes treated with SAT-CM or silenced

Purified cells were harvested from culture plates using 5 mM Na-EDTA (PanReac AppliChem) in PBS, pH 7.5 and incubated for 15 minutes at RT with Human TruStain FcX (Biolegend) to block Fc receptors. Depending on the experiment, cells were stained for 15 minutes with combinations of the following antibodies: monoclonal antibody PE-conjugated anti-CD300e (clone UPH2, Biolegend, 339704), monoclonal antibody BV421-conjugated anti-CD36 (clone 5-271, Biolegend, 336229), monoclonal antibody APC-conjugated anti-phospho AKT1 (S473, clone SDNRN, Thermo Fisher Scientific, 17971542). For phospho AKT1 labeling cells were fixed with 3.6% formaldehyde and permeabilized with PermWash solution (1% FBS, 0.2% saponin in PBS). The BD Horizon™ fixable viability stain 780 (BD Biosciences) was used to exclude dead cells from the analysis. Cells were washed, resuspended in FACS buffer and analysed by LSR Fortessa X-20 Cell Analyzer (BD Biosciences). Data were analysed using FlowJo software.

#### Mouse SVF

0.5 × 10^6^ cells isolated from SAT and VAT stromal vascular fraction, as described above, were incubated with TruStain FcX PLUS (BioLegend) for 10 min at 4 °C to block Fc receptors. Depending on the experiment, cells were stained with combinations of the following antibodies: PE-Cy7-conjugated anti-CD45 (clone 30-F11, Biolegend, 103114), BV785-conjugated anti-CD11b (clone M1/70, Biolegend, 101243), BV421-conjugated anti-F4/80 (clone BM8, Biolegend, 123132), BV650-conjugated anti-CD4 (clone RM4-5, Biolegend, 100546), AlexaFluor700-conjugated anti-mouse CD8a (clone QA17A07, Biolegend, 155022).

The BD Horizon™ fixable viability stain 780 (BD Biosciences) or Zombie Aqua Fixable Viability Dye (Biolegend) were used to exclude dead cells from the analysis. Cells were resuspended in FACS buffer and analysed by LSR Fortessa X-20 Cell Analyzer (BD Biosciences). Data were analysed using FlowJo software.

#### Mouse peripheral blood

Peripheral blood was transferred in a round-bottom tube and incubated with lysis buffer (BD Pharm Lyse; BD Biosciences, Cat# 555899) to lyse erythrocytes. Cells were incubated for 10 minutes at 4 °C with TruStain FcX PLUS (BioLegend) and stained with the monoclonal antibodies FITC-conjugated anti-CD45 (clone 30-F11; BioLegend, 103108), PE-conjugated anti-CD19 (clone 1D3/CD19; BioLegend, 152408), PE-Cy7-conjugated anti-Ly6G (clone 1A8; BioLegend, 127618), PerCP/Cyanine5.5-conjugated anti-CD8a (clone 53-6.7; BioLegend, 100734), APC-conjugated anti-CD4 (clone RM4-5; BioLegend, 100516), APC-Cy7-conjugated anti-CD3e (clone 145-2C11; BioLegend, 100330), BV421-conjugated anti-Ly6C (clone HK1.4; BioLegend, 128032), and BV785-conjugated anti-CD11b (clone M1/70; BioLegend, 101243). Zombie Aqua Fixable Viability Dye was used to exclude dead cells. Cells were resuspended in FACS buffer and analyzed by LSR Fortessa X-20 Cell Analyzer (BD Biosciences). Data were analysed using FlowJo software.

#### MME cells

Cells were harvested from culture plates using 5 mM Na-EDTA (PanReac AppliChem) in PBS, pH 7.5 and incubated for 15 minutes at RT with Human TruStain FcX (Biolegend) to block Fc receptors. Cells were stained for 15 minutes with monoclonal antibody PE-conjugated anti-CD36 (clone HM36, Biolegend, 102606), fixed with 3.6% formaldehyde, permeabilized with PermWash solution (1% FBS, 0.2% saponin in PBS), and labelled with monoclonal antibody APC-conjugated anti-phospho AKT1 (S473, clone SDNRN, Thermo Fisher Scientific, 17971542). The BD Horizon™ fixable viability stain 780 (BD Biosciences) was used to exclude dead cells from the analysis. Cells were washed, resuspended in FACS buffer and analysed by LSR Fortessa X-20 Cell Analyzer (BD Biosciences). Data were analysed using FlowJo software.

### RNA extraction and qRT-PCR

Snap-frozen tissue pieces were disrupted in TRIzol reagent (Thermo Fisher Scientific) using TissueLyser II (QIAGEN), while cell pellets were resuspended in TRIzol and disaggregated by pipetting, and RNA was extracted following manufacturer instructions. After quantification using NanoDrop Lite Spectrophotometer (Thermo Fisher Scientific), 1 μg of RNA was retrotranscribed using the High-Capacity cDNA Reverse Transcription Kit (Thermo Fisher Scientific) and the retrotranscription reaction conditions recommended by the manufacturer. 5 ng of cDNA were used in qRT-PCR reaction performed in QuantStudio™ 5 Real-Time PCR System (Applied Biosystems, Thermo Fisher Scientific); the reaction was performed using PowerUp™ SYBR™ Green Master Mix for qPCR (Applied Biosystems). For human samples, data were normalized to the endogenous reference gene β-actin; for mouse samples, data were normalized to the endogenous reference gene 18S and confirmed using a different reference gene (β-actin; data not shown). Primer sequences are reported in Supplementary Table S4.

### Sample preparation for MS-based proteomics

Cell pellets were resuspended in 30 μl of LYSE buffer (PreOmics GmbH, Planegg, Germany), containing reducing and alkylating reagents. After heating at 95 °C for 5 minutes, samples were subjected to mechanical disruption for 10 minutes using a BeatBox apparatus (PreOmics) and further sonicated in a Bioruptor (Diagenode, Seraing, Belgium) for 15 minutes at a 50% duty cycle. Protein concentration in the lysate was estimated based on tryptophan fluorescence compared to a standard curve^47^. For each sample, 2 ug of lysate were digested with 0.25 μg of trypsin and 0.25 μg LysC overnight at 37 °C under continuous shaking. The lysate was desalted and loaded onto StageTip plugs of SDB-RPS. Purified peptides were eluted with 80% acetonitrile-1% ammonia and dried, then further loaded onto EvoTips (Evosep Biosystems, Odense, Denmark).

### Liquid chromatography and mass spectrometry analysis

Samples (150 ng of purified peptides) were analyzed in an Orbitrap Astral mass spectrometer (Thermo Fisher Scientific) coupled to an EvoSep chromatography system (EvoSep Biosystems), using a commercial analytical column (IonOpticks, Melbourne, VIC, Australia) and a 60-samples-per-day method, with 21 minutes per run protocol. The mass spectrometer was operated in a data-independent acquisition (DIA) mode, consisting of one MS1 full scan (380–980 m/z range) acquired at a resolution of 240,000 and MS2 scans collected with isolation windows of 2 m/z, 300 scan events at HCD collision energy of 25%, with a scan range of 150-2000m/z. Raw files were quantified with the DIA-NN software (version 1.8.1)^48^ using a direct-DIA approach. Our proteomic analysis quantified 9290 protein groups in total, with an average of 7001 per sample. Pearson correlation ranged between 0.71 and 0.91 for different samples pairs and was 0.89 on average in the dataset.

### Differential protein abundance analysis

The raw files generated above were subjected to the standard “proDA” (v1.20.0) pipeline using R (v4.4.3). Only proteins belonging to the list in Supplementary Table S1 were considered as mapping in cellular processes involved in lipid and glucose metabolism and energy production.

Briefly, the abundance matrix was log2 converted and normalized using the “median_normalization” function with standard parameters. We than fit the probabilistic dropout model with “proDA” function considering in the design only the genotype, or the interaction of the genotyope and the diet. Finally, we identify differentially abundant proteins with the “test_diff” function. As proteomics data were used as an orthogonal validation of findings obtained from independent experimental assays, we used unadjusted p values (p < 0.05), without correction for multiple testing, due to limited statistical power associated with the targeted analysis and sample size.

The resulting enriched abundance protein were subjected to gene ontology analysis using the R packages “clusterProfiler” ^49^ (version 4.14.6) and “msigdbr” ^50,51^ using the categories H and C2 for *mus musculus*.

### Oxygen consumption rate (OCR) measurements

For OCR measurements, BMDM were seeded at a density of 5× 10^4^ cells per well in XF24-well cell culture microplates (Agilent Technologies, Santa Clara, CA, USA) in MMe polarization medium. After 24 hours, the culture medium was replaced with DMEM, supplemented with 10 mM glucose and 5 mM HEPES. Thirty minutes later, OCR measurement was measured in real time using an XF24 Extracellular Flux Analyzer (Agilent Technologies). A preliminary titration was performed to identify the concentration of carbonyl cyanide-p-trifluoromethoxyphenylhydrazone (CCCP, 1.5 μM) that maximally increased OCR. OCR was measured under basal condition and following sequential addition of 2 μM oligomycin to assess ATP-linked respiration, 1.5 μM CCCP to assess maximal respiration, and 1 μM rotenone plus 1μM antimycin A to assess non-mitochondrial respiration. OCR values were normalized for total protein content.

### Fatty acid and glucose uptake assay

To assess the ability of MMe and silenced human monocytes to uptake fatty acids or glucose, two fluorescent analogues were used: BODIPY FL C16 (Thermo Fisher Scientific) and 2-NBDG (Thermo Fisher Scientific), respectively. Cells were incubated at 37 °C for 30 minutes with 5 µg/ml BODIPY FL C16 to evaluate fatty acid uptake, or with 50 µM 2-NBDG to assess glucose uptake. After incubation, cells were washed twice with PBS, detached, and analyzed by flow cytometry.

### Statistical analysis

Data analysis and graphing were performed using GraphPad Prism 8.02 (GraphPad Software, San Diego, CA, USA). For comparisons between two conditions, a two-tailed paired Student’s t-test was used. For comparisons involving three or more conditions with one or two independent variables, one-way or two-way analysis of variance (ANOVA) was employed, respectively, followed by Bonferroni post hoc correction. P values are defined as *p < 0.05, **p < 0.01, ***p < 0.001.

## Abbreviations

AdExos: Adipocyte-Derived Exosomes
AT: Adipose Tissue
ATM: Adipose Tissue Macrophages
BMI: Body Mass Index
CM: Conditioned Medium
CFD: Control-Fat Diet
CLS: Crown-Like Structures
FDR: False Discovery Rate
FBS: Fetal Bovine Serum
FFA: Free Fatty Acids
GTT: Glucose Tolerance Test
H&E: Hematoxylin and Eosin
HFD: High-Fat Diet
ITAM: Immunoreceptor Tyrosine-based Activation Motif
ITT: Insulin Tolerance Test
i.p.: Intraperitoneal
KO: Knock-Out
LAM: Lipid-Associated Macrophages
MFI: Mean Fluorescence Intensity
MASLD: Metabolic Dysfunction-Associated Steatotic Liver Disease
MMe: Metabolically Activated Macrophages
SVF: Stromal Vascular Fraction
SAT: Subcutaneous Adipose Tissue
TAG: Triacylglycerols
VAT: Visceral Adipose Tissue
WAT: White Adipose Tissue
WT: Wild-Type

## Ethics approval and consent to participate

Peripheral blood and SAT were collected from lean and obese participants in accordance with the principles of the Declaration of Helsinki. Written informed consent was obtained all participants prior to enrolment, in accordance with the study protocol approved by the “Comitato Etico Territoriale Area Centro – Est Veneto (CET – ACEV)” (protocol number: 2892P).

All animal experiments were conducted in accordance with national guidelines for animal welfare and approved by the Italian Ministry of Health (protocol number: 858/2023-PR).

## Availability of data and materials

The mass spectrometry proteomics data have been deposited to the ProteomeXchange Consortium via the PRIDE^52^ partner repository with the dataset identifier PXD072156.

Any additional information required to reanalyze the data reported in current study are available from the corresponding author upon reasonable request.

## Authors’ contributions

SC, MdB, AG, and RV conceived the study. SC, SP, SV, ET, SG, AB, and ADM performed the experiments. FC and MM performed the lipidomic and proteomic analyses, respectively. GS and ML performed the data analysis. SC and MdB supervised the experiments and drafted the manuscript. SS and CM provided intellectual input on data presentation and manuscript structure. All authors read and approved the final manuscript.

## Acknowledgements

We are grateful to Professor G. Codolo from the University of Padua for kindly providing the CD300e constitutive KO (Cd300e^−/−^) and Cd300e^LysM^ mice from C57BL/6J background. We thank Prof. Matthias Mann for providing access to mass spectrometry instrumentation at the Max Planck Institute of Biochemistry (Martinsried, Germany) for the opportunity to perform MS-based proteomics in his laboratory.

