## Supplementary Figures and tables for "CD300e modulates metabolic programs in adipose tissue macrophages during obesity"

**Supplementary Files:**

- Supplementary Figure 1
- Supplementary Figure 2
- Supplementary Figure 3
- Supplementary Table S2
- Supplementary Table S3
- Supplementary Table S4

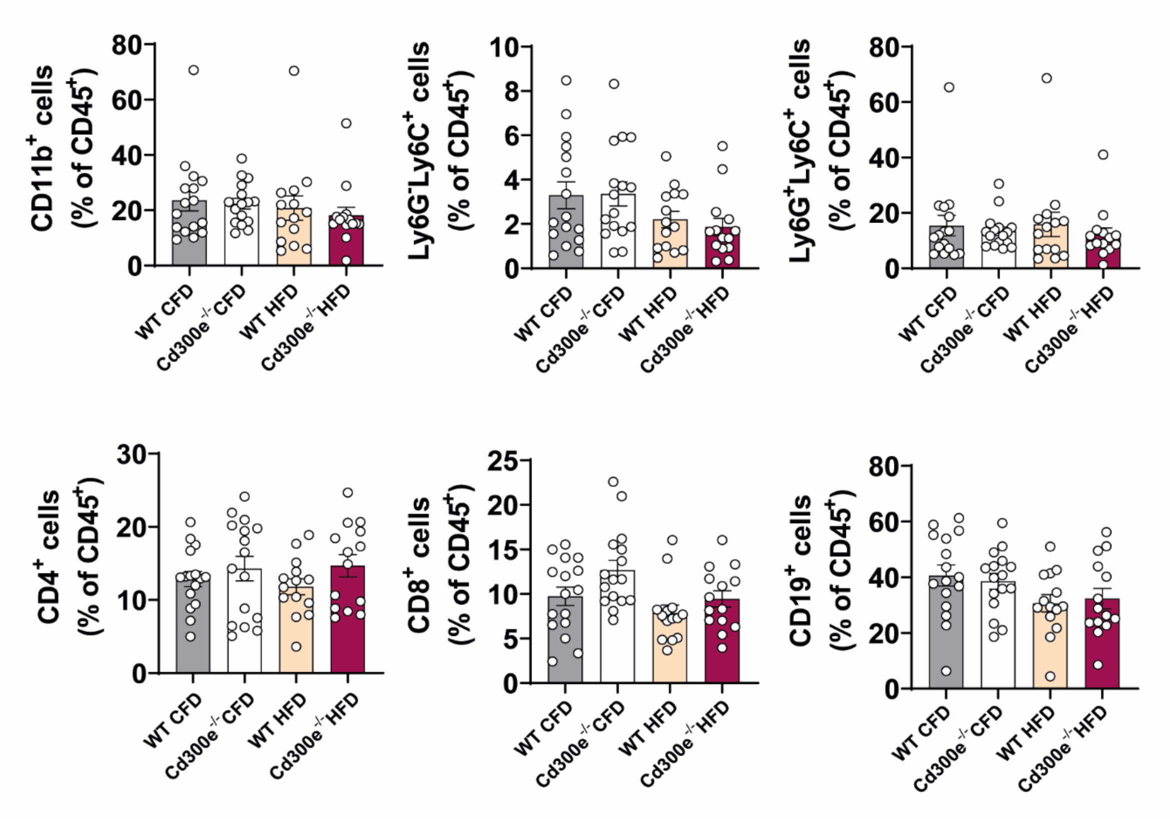


**Supplementary Figure 1. Blood immune cell composition are not influenced by the diet regimen and the absence of CD300e.** The percentage of CD11b$^+$ myeloid cells, Ly6G$^-$Ly6C$^+$ monocytes and Ly6G$^+$Ly6C$^+$ neutrophils, CD4$^+$ and CD8$^+$ T lymphocytes, and CD19$^+$ B lymphocytes were determined within total blood CD45$^+$ leukocytes collected from WT and *Cd300e*$^{-/-}$ mice after 16 weeks of CFD or HFD feeding. Data are expressed as percentage (%) of CD45$^+$ cells (mean ± SEM of n=16/genotype of CFD-fed mice and n=14/genotype of HFD-fed mice).

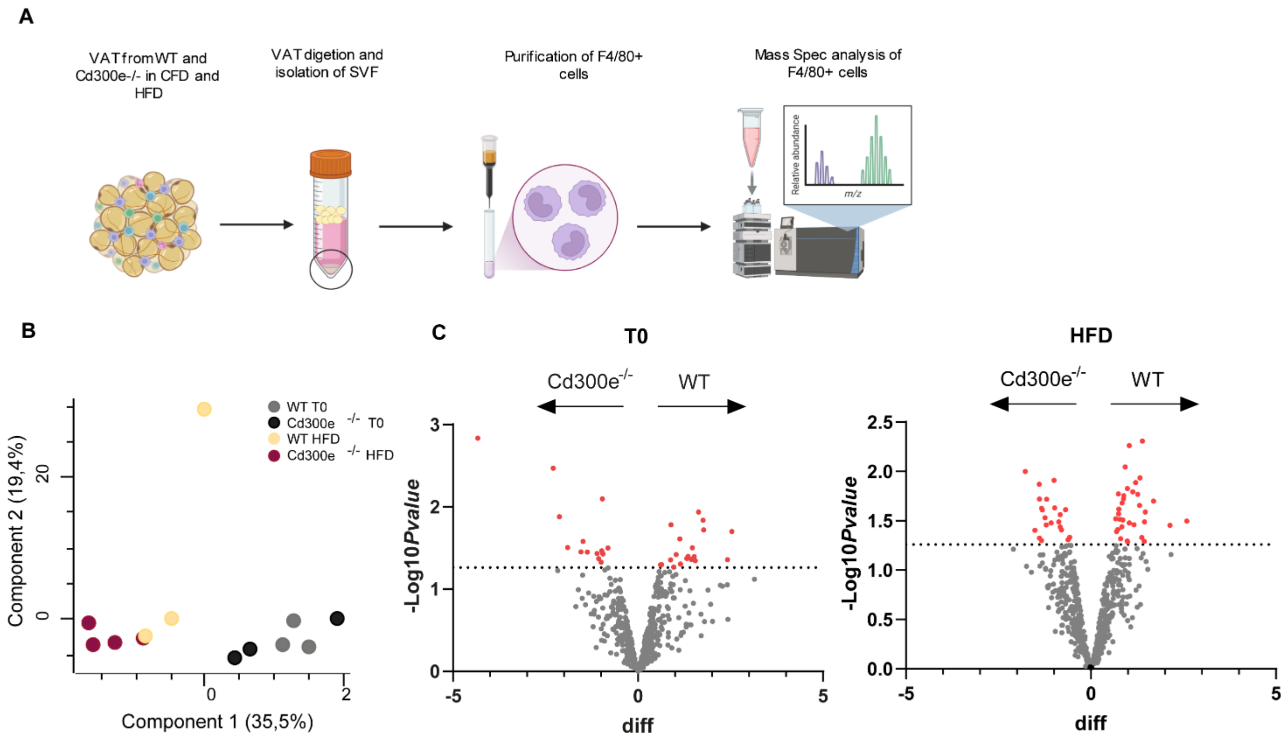


**Supplementary Figure 2. Proteomic analysis of *Cd300e*$^{-/-}$ ATMs.**
(A) Schematic overview of the mass-spectrometry-based proteomic workflow. (B) PCA analysis of the proteomic data. (C) Volcano plot showing enriched proteins between genotypes at $T_0$ (left panel) or under HFD (right panel).

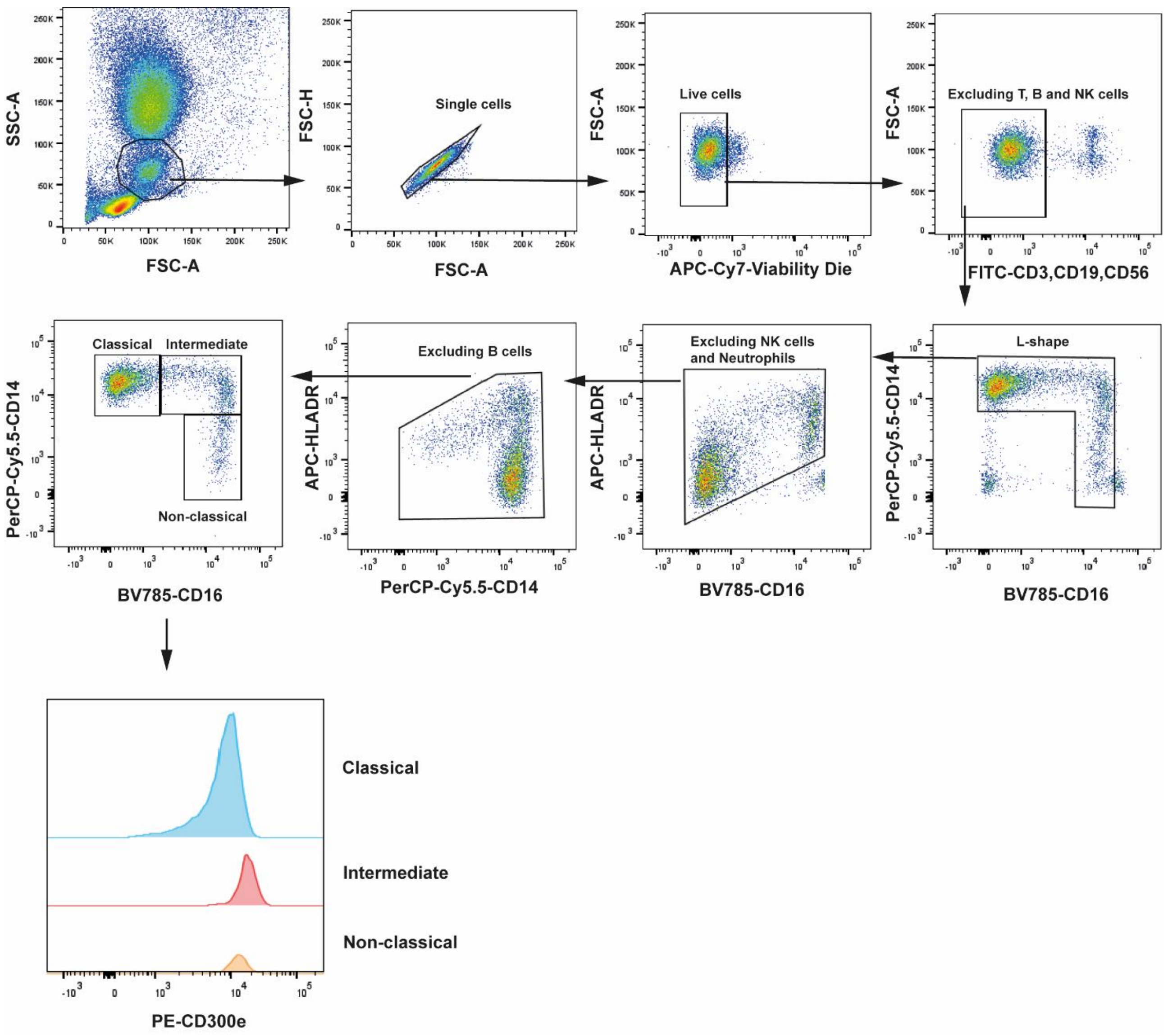


**Supplementary Figure 3. Gating strategy for the identification of the circulating monocytes subsets.** Forward and side scatter were used to isolate the interesting population. Subsequently T, B and NK cells were excluded and the CD3$^-$, CD19$^-$ and CD56$^-$ cells identified within the single cell and live cell gates. Monocytes were identified and divided into classical (CD14$^+$, CD16$^-$), intermediate (CD14$^+$, CD16$^+$) and non-classical monocytes (CD14$^-$, CD16$^+$). Then, any further contamination of NK cells, neutrophils (CD16$^+$, HLA-DR$^-$) and B cells (HLA-DR$^+$, CD14$^-$) has been excluded and within the three subclasses of monocytes the fluorescence intensity of CD300e has been analyzed.

| | **Lean** | **Obese** |
|---|---|---|
| **Sex** | **18 F and 17 M** | **24 F and 19 M** |
| **Age (mean ± SEM)** | **(34.6 ± 1.6)** | **(42.9 ± 1.4)** |
| **Height (mean ± SEM)** | **(172.8 ± 1.8)** | **(169.8 ±1.4)** |
| **Weight (mean ± SEM)** | **(64.8 ±2)** | **(122.5 ± 3.5)** |
| **BMI (mean ± SEM)** | **(21.6 ±0.3)** | **(42.5±1.1)** |

**Supplementary Table S2. Clinical characteristics of lean and obese patients.**

| **siRNA** | **Sequence** |
|---|---|
| *Hs_IREM2_1* | TCCCATCTTCCTGGTGGTGAA |
| *Hs_IREM2_2* | CAGAACCTCAATGAAGATGAT |
| *Hs_IREM2_3* | GAGCATGTACAAGGGATATAA |
| *Hs_IREM2_5* | CAGTGTGGTGTCAGTATGAGA |

**Supplementary Table S3. SiRNA sequence used for CD300e silencing.**

| Primer | Sequence 5' – 3' |
|---|---|
| h β-actin FW | TGAGATGCGTTGTTACAGGA |
| h β-actin RV | ACGAAAGCAATGCTATCA |
| h CD300e FW | GGGAGGTGTTGACCCAAAAT |
| h CD300e RV | AGGACCACGAGCAGGAAGT |
| m β-actin FW | GATTACTGCTCTGGCTCCTAGC |
| m β-actin RV | GACTCATCGTACTCCTGCTTGC |
| m 18S FW | TGTCTCAAAGATTAAGCCATGC |
| m 18S RV | GCGACCAAAGGAACCATAAC |
| m CD300e FW | CCAGTGAGGCAGGAGGATGAG |
| m CD300e RV | AGAGACAAACAACCTTGGAAGCAGAG |
| m IL-6 FW | TAATTCATATCTTCAACCAA |
| m IL-6 RV | TCCTTAGCCACTCCTTCTGT |
| m TNF-α FW | ACTGAACTTCGGGGTGAT |
| m TNF-α RV | CTGAGTGTGAGGGTCTGG |
| m IL-1β FW | CTGCTTCCAAACCTTTGACC |
| m IL-1β RV | AGCTTCTCCACAGCCACAAT |
| m ACC FW | GGCCAGTGCTATGCTGAGAT |
| m ACC RV | AGGGTCAAGTGCTGCTCCA |
| m ATGL FW | CAGCACATTTATCCCGGTGTAC |
| m ATGL RV | AAATGCC GCCATCCACATAG |
| m SREBP1 FW | GCGCTACCGGTCTTCTATCA |
| m SREBP1 RV | TGCTGCCAAAAGACAAGGG |
| m ChREBP FW | AAGTCCTTGGTCGGGAAGTATACA |
| m ChREBP RV | ACTCCCTCAAAGTCATCACAAACA |
| m FAS FW | CTGCGGAAACTTCAGGAAATG |
| m FAS RV | GGTTCGGAATGCTATCCAGG |

**Supplementary Table S4. Primers sequence used for qRT-PCR.**
